# A lab-on-a-chip model of glaucoma

**DOI:** 10.1101/704510

**Authors:** Fatemeh Nafian, Babak Kamali Doust Azad, Shahin Yazdani, Mohammad Javad Rasaee, Narsis Daftarian

**Author notes:** FN and BKDA has contributed equally to this work and should be considered as co-first authors. MJR and ND are co-corresponding authors. All queries and request for reprints should be addressed to MJR.

## Abstract

We developed a glaucoma-on-a-chip (GOC) model to evaluate the viability of retinal ganglion cells (RGCs) against high pressure and the potential effect of neuroprotection. A three-layered chip consisting of interconnecting microchannels and culture wells was designed based on simulation of physical parameters. The chip layers were fabricated from poly-methyl methacrylate sheets. Multiple inlet ports allow culture media and gas into the wells under elevated hydrostatic pressure (EHP). The bottom surface of the wells was modified by air plasma and coated with different membranes to model an extracellular microenvironment. The SH-SY5Y neuroblastoma cell line served as model cells to determine the best supporting membrane which was revealed to be PDL/laminin. Thereafter, the study experiments were performed using RGCs obtained from postnatal 5-7 Wistar rats purified by magnetic assisted cell sorting. Flow cytometry and immunocytochemistry assays demonstrated 70% purification for RGCs. The cultured RGCs were exposed to normal (15 mmHg) or elevated pressure (33 mmHg) for 6, 12, 24, 36 and 48 hours, with and without adding brain-derived neurotrophic factor (BDNF) or a novel BDNF mimetic (RNYK). RGC survival rates were 85, 78, 70, 67 and 61 percent under normal pressure versus 40, 22, 18, 12 and 10 percent under high pressure at 6, 12, 24, 36 and 48 hours, respectively (P <0.0001). BDNF and RNYK treatments induced separately an approximate two-fold decrease in the rate of RGC death under both normal and elevated pressures (p <0.01 to 0.0001). This GOC model recapitulated the effects of elevated pressure during relatively short time periods and demonstrated the neuroprotective effects of BDNF and RNYK.

## 1. Introduction

Glaucoma is a collective term used to define a group of neurodegenerative processes affecting the entire visual pathway best distinguished by progressive, irreversible destruction and death of retinal ganglion cells (RGCs). The disease spectrum is estimated to affect more than 100 million people worldwide by the year of 2040 [1]. The primary risk factor for progression and development of glaucoma is elevated intraocular pressure (IOP). IOP is regulated by the balance between aqueous humor secretion into the anterior chamber on one hand and its drainage via the trabecular meshwork (conventional outflow) or the uveoscleral outflow pathways on the other hand [2]. The average value for normal IOP in most population based human studies is 14-17 mmHg with 95% confidence intervals of 10-21 mmHg [3].

Several experimental in vivo glaucoma models have been developed and used for studies on glaucoma mechanisms. These include laser photocoagulation of the Perilimbal region [4], red blood cell or microbeads injection into the anterior chamber [5, 6], Episcleral vein obstruction [7], and Episcleral vein saline injection [8–10]. Without sequential treatments, the duration of IOP elevation is transient in these models. Precise control over IOP elevation is difficult and certain problems may occur; these include intraocular inflammation, irreversible mydriasis, IOP variability, hyphema, reduced visibility of optic discs, corneal opacity, and scleral burns [11, 12]. While in vivo animal models are indispensable to determine what events occur in live organisms, these strategies typically involve poorly defined and uncontrollable factors and the results can be difficult to understand at cellular and molecular levels [13].

Ishikawa et al. designed an ex vivo hydrostatic pressure model which demonstrated better retention of neuron-neuron and neuron-glial interactions in dissected eye cups [14–18]. This model excluded the effects of ischemia and allowed studying of the direct effects of pressure on the retina. Eyecups were sunken to the bottom of a glass cylinders containing different liquid heights. Although ex vivo systems produce reliable results, there are several limitations. For example, the incubation period is limited by the duration of tissue viability because survival factors originating from axonal transport or the bloodstream are eliminated in ex vivo models[13].

To overcome these limitations, in vitro models have been developed using cultured cells in supporting environments to clarify glaucoma mechanisms and appear as valuable tools to evaluate the response of individual cell populations against noxious conditions and novel treatments [19–21]. These models have studied optic nerve head astrocytes, RGCs and other types of retinal cells utilizing pressure loading systems [21–28]. Elevation of pressure constitutes the gold standard to model ocular hypertension in vitro and has gained increasing attention in recent years. Elevated hydrostatic pressure (EHP) systems provide remarkable information on cell apoptosis, elastin synthesis, cell migration, and production of cell adhesion molecules. EHPs have provided new insights into the molecular and cellular mechanisms of glaucoma.

Recently, lab-on-a-chip (LOC) technology has been developed. This approach entails a simpler and smaller analysis platform, resulting in better temporal and spatial control of local cellular microenvironments, passive and active cell handling, less consumption of reagents, faster test results, economizing study logistics and energy savings [29–32]. With the goal of transferring costly laboratory equipment onto small, user-friendly, easily replicable chips, LOC technology has dramatically altered many fields such as medicine, biochemistry, and biotechnology.

The aim of the present study was to design and establish a glaucoma-on-a-chip (GOC) model consisting of an EHP system, coupled to a microculture system using purified primary rat RGCs. To achieve this aim, we designed pressure chips in which HP could be modulated with the added benefits of higher speed, greater precision and finer control than previous EHP systems. We studied RGC survival under normal versus elevated hydrostatic pressures and also compared survival of these cells when treated with a neuroprotective growth factor (brain-derived neurotrophic factor, BDNF) or a mimicry peptide, named RNYK, as a putative agonist of neurotrophic tyrosine kinase receptor type 2 (NTRK2) [33]. BDNF is a well-known neuroprotectant and a potent therapeutic candidate for neurodegenerative diseases. However, there are several clinical concerns about its therapeutic applications. RNYK as a mimicking small peptide (similar to loop2 of BDNF) is an alternative to circumvent these problems. The RNYK neuroprotective effect has been confirmed with equal efficacy to or even better than BDNF.

## 2. Material and methods

### 2.1. Design of a three-layered chip

In order to induce a high pressure environment to simulate glaucomatous conditions, several three-layered EHP models were designed (Fig. 1). A number of morphological parameters were considered including number and location of feeding ports in the top layer (layer A), the shape of micro-channels and wells in the middle layer (layer B), and surface modification for the bottom layer (layer C) of the chip. AutoCAD 2014 was used to design the layers. Some physical parameters including gas concentrations and fluid velocity profile for media at seeding/feeding time were simulated by Comsol Multiphysics 5.2a software employing finite element methods. Two-dimensional incompressible Navier-Stokes equations were used along with the continuity equation to determine the liquid velocity profile of culture media throughout the fluid dynamic field at the seeding and feeding time

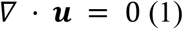

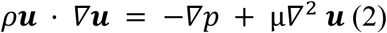

**Figure 1.**
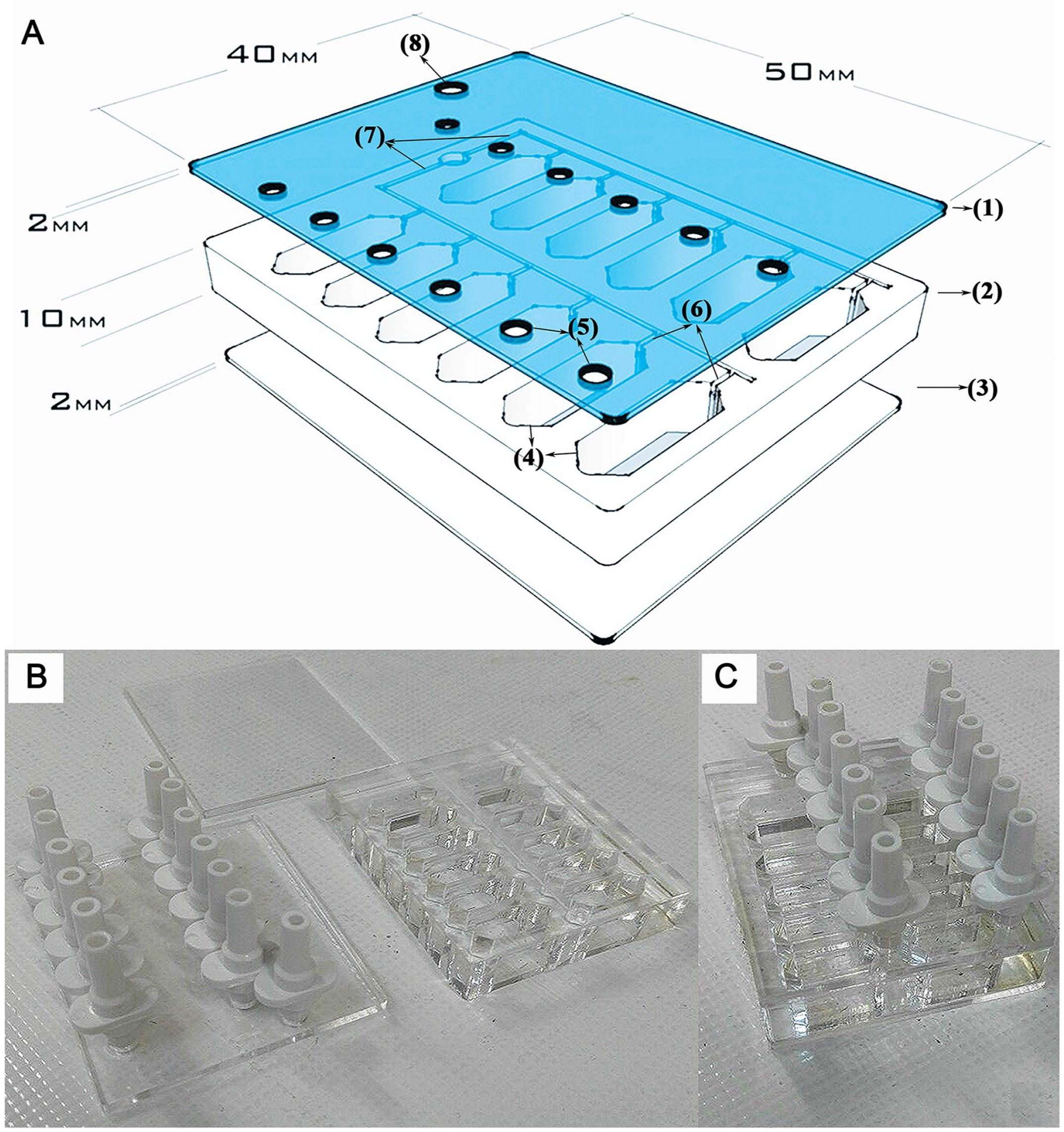
Schematic of a designed chip in AutoCAD software. (A) Three-dimensional design of layers A (1), B (2), and C (3) containing 12 wells (4), feeding ports (5), and microchannels (6), 2 main channels (7) and 1 single gas inlet (8). Layer A consisted of one feeding port per well and one main gas inlet. Layer B involved 12 hexagonal wells and microchannels. Layer C provided a surface area for cell cultures. Layers were fabricated (B) and assembled (C) to establish a complete chip.

Where **u** is the velocity vector, p is the pressure of injection, *ρ* is the density of media, and μ is the dynamic viscosity. The no-slip boundary condition was assumed at the walls. Constant velocity was specified at the inlet. The outlet was assumed to be at atmospheric pressure.

After the flow reached a steady-state condition, gas was assumed to be transported only by diffusion so that gas distribution was estimated using Fick’s law (FL). Gas was released at the inlet at a specific concentration (C_0_), which was monitored after a certain sampling time period. The transient two-dimensional mass transport equation is as follows:

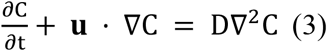

Where C is the gas concentration in the bulk, and D is the diffusion coefficient.

A novel low-cost fabrication method was established to produce the three-layered chip entirely from poly-methyl methacrylate (PMMA, Cho Chen, Company, city, country). PMMA is the most common member of the acrylic family. The optical clarity and ability to resist environmental stress make PMMA ideal for replacing glass in light transmission applications [34]. PMMA biocompatibility has been established through years in medicine and ophthalmology [35].

### 2.2. Fabrication and testing of the three-layered chip

A laser system (fiber laser, PNC laser Co. Ltd, Tehran, Iran) was used to cut polymerized PMMA sheets of about 10mm thickness for layer B and 2mm for layers A and C. The laser processing system was equipped with a programming system (NC system), an automatic programming combining computer and CAD technology. When using the laser for fabricating channels and wells on PMMA sheet, the laser power and scanning speed were considered as main factors. Several tests were performed to explore the correct correlation between laser power and scanning speed. A 25W laser power and a 300mm/s laser scanning speed were selected to produce main channels and sub-channels on layer B. In addition, 20W and 45W laser power were applied to cut PMMA sheets 10 mm and 2 mm in thickness respectively, at 100 mm/s laser scanning speed to make the wells and ports in layers A and B.

A thermal-solvent method was used to attach the three layers of any single chip. This method is based on dissolving PMMA in a thin layer of a solvent (isopropyl alcohol) between two PMMA substrates at 75 °C for 10 min [36]. Adhesion of PMMA substrates involved the following steps. First, the layers cut by the laser were rinsed in deionized water for 5 minutes to remove tiny debris and dried at 40 °C for 10 minutes in an oven. Next the solvent was added between layers B and C which were aligned and fixed using paper clamps to avoid entrapment of air bubbles between the binding surfaces. The attached sheets were heated and finally treated with air plasma. Radiofrequency discharged plasma was generated by a dinner plasma generator from Dinner CO. at 13.6 MHZ frequency and 20W power for 10 min. The base pressure of the plasma reactor was increased to 0.35 mmHg after gas feeding. After attaching layers B and C, layer A was attached using the same method described above to complete the three-layered chip.

### 2.3. Extracellular microenvironment model on modified PMMA surface

SH-SY5Y cell line which is the third sequential sub-clone of the SKN-SH human neuroblastoma cell line was purchased from the Pasteur Institute (Tehran, Iran). These cells phenotypically change from neuronal type (N) to surface adherent type (S) by all-trans-retinoic acid (ATRA) treatment [37]. The cells were seeded at an initial density of 10^4^ cells per cm^2^ in the wells of the chip, which were pre-coated with one of the following membranes: collagen (0.05 mg ml^−1^), ploy-D-lysine (PDL)/laminin (0.01 mg ml^−1^/0.05 mg ml^−1^), gelatin (1 mg ml^−1^), and a composite nanofiber of polyacrylonitrile (PAN) and polyaniline (PANI), named PAN/PANI. When the cells were approximately 75% confluent, ATRA (Sigma-Aldrich, Vienna, Austria) treatment was performed at a final concentration of 10 μM in Dulbecco’s modified Eagle’s medium (DMEM) (Gibco-BRL *Life Technologies*, *Grand Island*, NY) supplemented with 2 mM L-glutamine, penicillin (20 U ml^−1^), streptomycin (20 mg ml^−1^), and 15% (by volume) Fetal Bovine Serum (*FBS*) (*Gibco-BRL Life Technologies*) and incubated for 5 days. Cell morphology, density, and distribution were compared among different coating materials.

### 2.4. Purification of RGCs and primary culture in the chip wells

Forty-eight hours before the experiment, chip surface was modified using plasma jet and exposed to UV light for 1h to assess successful sterilization. Then, 500 μl of PDL solution (Sigma-Aldrich), 10 μg ml^−1^ was added to each well for 1h. The wells were washed three times with sterile water and completely dried. The day before the experiment, 500 μl of laminin solution (5 μg ml^−1^, Sigma-Aldrich) prepared in Neurobasal medium was added to each well and incubated overnight at 37°C (8% CO2). The laminin/Neurobasal solution was replaced with 500 μl of “retinal ganglion cell media” which was prepared as follows: Neurobasal medium (Gibco-BRL Life Technologies) with 1% bovine serum albumin (BSA, Sigma-Aldrich), sodium selenite (6.7 ng ml^−1^, Sigma-Aldrich), Insulin-Transferrin-Selenium (ITS-G) (100X) (Thermo Fisher Scientific, Grand Island, New York, USA), progesterone (20 nM; Merck Millipore), B27 Supplement (50X) (Thermo Fisher Scientific), sodium pyruvate (1 mM; Merck Millipore), L-Glutamine (100x) (2 mM; Gibco-BRL Life Technologies), penicillin (20 U ml^−1^), and streptomycin (20 mg ml^−1^).

In accordance with the regulations of association for research in vision and ophthalmology (ARVO) for animal studies, ten postnatal day’s five to seven Wistar rats (Razi Institute, Karaj, Iran) were sacrificed by decapitation and then enucleated. The cornea, lens, and vitreous humor were removed and the retina was gently lifted away using a small spatula. The retinas were stored at room temperature in 4.5 ml Hank’s balanced salt solution, Ca2+ and Mg2+ free (HBSS-CMF, Sigma-Aldrich), pH 7.4. Then, 0.5 ml of trypsin (2.5%, Gibco-BRL *Life Technologies*) was added to the tube containing five retinas and incubated in a water bath at 37°C for 15 min where the tube was gently swirled every 5 min. The HBSS/trypsin solution was replaced by 5 ml of HBSS (containing 50 μl of DNase I 0.4%) and incubated at 37°C for 5 min. The retina was converted into a cell suspension by triturating and homogenizing the solution without introducing bubbles. The cells were centrifuged at 300g for 5 min and re-suspended in 1ml HBSS containing 0.5 mg ml^-1^ BSA. The cell suspension was filtered through a mesh filter (pore size 40 μm, BD Falcon, Franklin Lakes, NJ) to yield a single cell suspension.

For magnetic assisted cell sorting (MACS), a concentration of 10^7^ retinal cells per 90 μl of DBPS/BSA buffer (Dulbecco’s PBS containing 0.5 mg ml^−1^ BSA) was incubated with 10 μl of the RGC-specific marker CD90.1 microbeads (Miltenyi Biotec, GmbH, and Bergisch Gladbach, Germany) at room temperature for 30 min and then washed with 5 ml of DPBS/BSA buffer. The cell suspension was passed through an LS column to hold the micro bead-labelled cells (RGCs) in the magnetic field of a MidiMACS system (Miltenyi Biotec). The adherent RGCs were rinsed with 3 ml of DPBS/BSA buffer to deplete endothelial cells and microglia. The column was removed from the separator and held over a new 15-ml tube. The adhered cells were flushed out by firmly pushing the plunger into the column and counted.

### 2.5. Flow cytometry and immunocytochemistry of RGCs

In different isolation steps, cells were stained with mouse anti-CD90.1 antibody (Millipore, Bedford, MA) and goat anti-mouse IgG antibody FITC conjugated (Santa Cruz Biotechnology, Inc., Dallas, USA) as primary and secondary antibodies, respectively. The anti-CD90.1 antibody was added to the cell suspension at a concentration of 0.1 μg per 100 μl for 10^6^ cells and incubated at 4°C for 20 min. The cells were washed three times with PBS buffer and then 100 μl anti-mouse IgG antibody FITC conjugated was added (1:100 dilution) and incubated at room temperature for 30 min in the dark. After 3-times washing, the fluorescence emission intensity was measured using a flow cytometer. FlowJo (Tree Star, Inc., Ashland, OR) was used to exclude dead cells from the analysis based on scatter signals. The number of cells exhibiting CD90.1 were manually counted, and the RGC percentage was calculated by determining the rate of CD90.1-positive cells relative to other ones.

Purified RGCs were cultured on a chip pre-coated with PDL/laminin at 37°C and 8% CO2. After a 3-day culture, RGCs were fixed with cold methanol (−10°C) for 10 min, dried, and then blocked with 300μl of PBS containing 1% BSA for 20 min at room temperature. The cells were incubated with 10 μl mouse anti-CD90.1 antibody (1:100 dilution) as a primary antibody overnight at 4 °C. RGCs were exposed to goat anti-mouse IgG antibody FITC conjugated as a corresponding fluorescent secondary antibody (1:100 dilution) at room temperature for 1 h in the dark. Nuclei were counterstained with 4’,6-diamidino-2-phenylindole (DAPI, Thermo Fisher Scientific, Pennsylvania, USA) as a colouring reagent. To determine the expression of specific marker on the RGC surface (CD90.1-positive cells), several random fields were imaged under a fluorescence microscope.

### 2.6. Measurement of Cell Viability

RGCs were seeded at an initial density of 10^4^ cells per cm2 in two sets of 12-well chips prior to the pressure experiment. RGCs were exposed to high hydrostatic pressures in a time-dependent manner for 6, 12, 24, 36 and 48 hours. BDNF (50 ng ml^−1^; Thermo Fisher Scientific, Waltham, USA) and RNYK peptide (5 ng ml^−1^) were separately applied to RGCs in one chip under normal and the other elevated hydrostatic pressure. RGCs in control groups did not receive BDNF/RNYK in the same experiments. The chip pressure was checked periodically by tonometer for fluctuations to hold the pressure level constant in each test. Cell viability was measured using 3-(4, 5-dimethylthiazol-2-yl)-2, 5-diphenyltetrazolium bromide (MTT, Sigma-Aldrich) assay for each well in the six groups. MTT solution was added to the media at a final concentration of 0.5 mg ml^-1^ and aspirated from the wells after 4 h. The residual crystals (formazan) were dissolved in dimethyl sulfoxide (DMSO, Sigma-Aldrich) and the absorbance was measured at near 570 nm. These experiments were repeated for at least three times to check the reproducibility of biologic effects. All measured variables were described as mean ±SD.

### 2.7. Statistical Analysis

Cell viability in BDNF/RNYK treatment groups was compared to that of untreated controls under normal and elevated pressure. The GraphPad Prism 7.0 statistical software was employed (San Diego, USA). Significance was determined at p <0.05 level using two-way ANOVA with Holm-Sidak’s multiple comparison test.

## 3. Results

### 3.1. Testing of the three-layered chip

The three-layered chip was designed to allow inflow of cell culture media from feeding inlets in layer A into the wells in layer B adjacent to layer C which acts the culture surface at the bottom of the wells (Fig. 1). We fabricated stand-alone feeding inlets for each well to let treat culture cells within the wells in an individual manner. We initially devised two feeding ports per well in layer A (the results not shown). The drawback to this design was inappropriate surface tension leading to an uneven distribution of the culture media. We then shifted to a single feeding port at one extreme of the well and placed a gas inlet on the other extreme to allow uniform liquid distribution. Locating the feeding port at the extreme end of the well reduced shearing stress and avoided cellular detachment and necrosis in the mid-region of the well.

Based on simulation results, a hexagonal (divergent-convergent) well performed better than a rectangular well. If the cells are seeded into a rectangular well, they mainly localize in the central zone due to maximum velocity there influenced by boundary layers related to sidewalls (Fig. 2A, red curve). Whereas, velocity profile became more even in a divergent-convergent input due to the reduction cross section area and mitigation of boundary layer effects (Fig. 2B, blue curve), resulting in a homogeneous flow velocity and cell distribution across the well at the feeding and seeding times, respectively. We designed 12 wells in each chip for simultaneously RGCs treatment with different compounds per well without any mixing of culture media or cells from one well to another via the gas channels during feeding process. The gas channels necessarily had to be interconnected to provide the same level of pressure and gas conditions to all wells within a chip. A single gas inlet was located for each chip which led to two separate main channels with six branching sub-channels (Fig. 2C).

**Figure 2.**
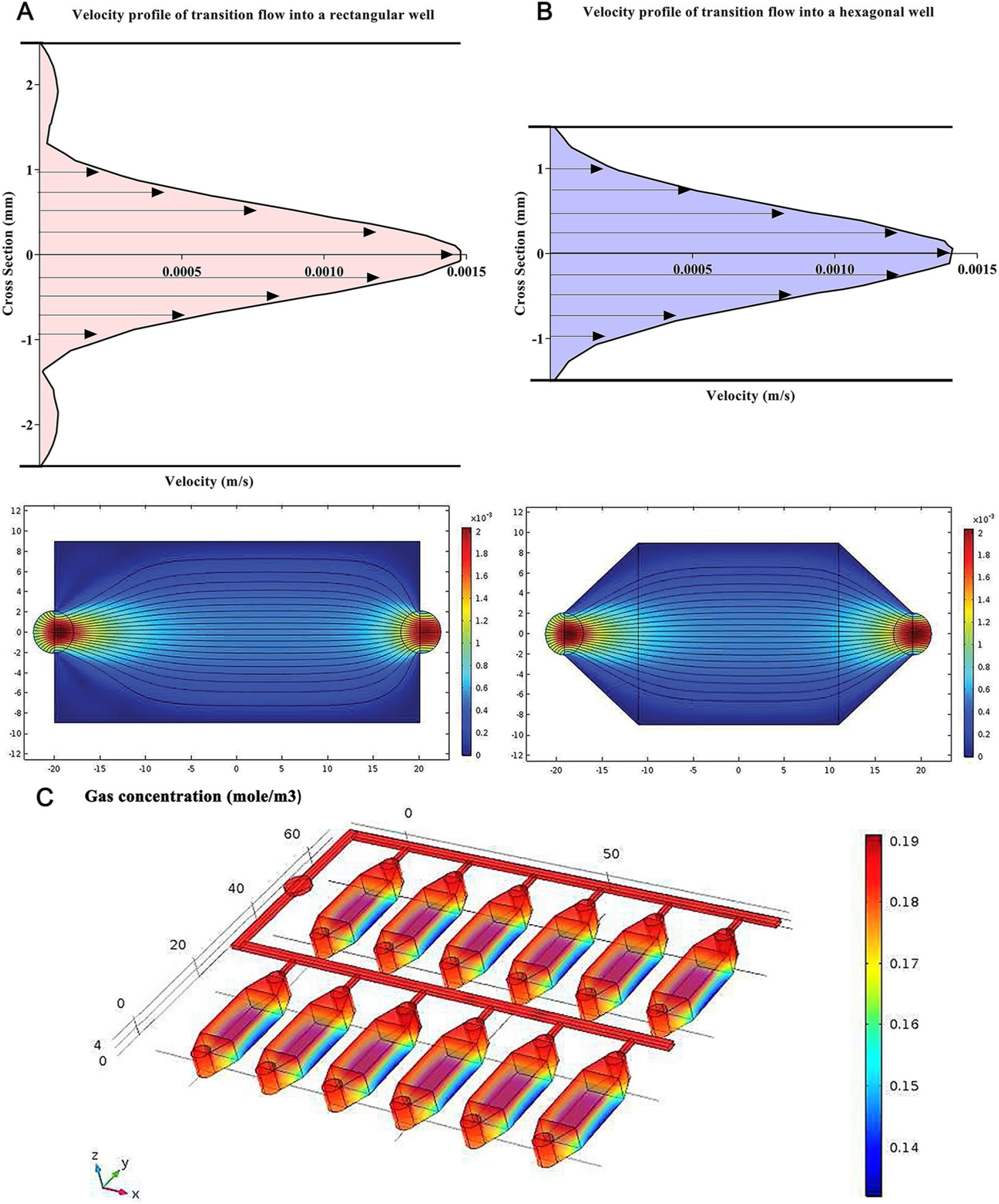
Estimated velocity profiles and gas concentration by Comsol Multiphysics software. (A) An entrance region was observed where the upstream flow entered the rectangular well. The boundary layer increased downstream, delayed the axial flow at the wall and accelerated the core flow in the center by maintaining the same flow rate. (B) The velocity profile became more even across the hexagonal well by straitening the walls at the entrance region and reducing the boundary layers at downstream (C) Gas concentration slightly changed throughout the main channels, sub-channels, and wells.

The bottom surface of the wells (layer C) were treated by air plasma to chemical modification and functionalization. This surface treatment increases surface roughness, amount of oxygen atoms, and functional hydrophilic groups (and as a result the number of polar interactions), especially the electron-donor parameters. These modifications enhance cellular adhesion which is fundamental for cell growth, migration, and differentiation.

### 3.2. Extracellular microenvironment model on modified PMMA surface

ATRA was added to the culture media to differentiate SH-SY5Y neuroblastoma cells into more mature, neuron-like cells. ATRA is a powerful growth inhibitor but promotes normal cellular differentiation. This low cost procedure is easily performed as compared to primary neuronal culture and may generate a homogenous neuronal cell population. The morphology and distribution of neuron-like cells were distinguished by phase-contrast microscopy (Fig. 3A). Untreated neuroblastoma cells possess large and flat cell bodies, however neuron-like cells have branching neuronal networks with smaller and round cell bodies. The neuron-like cells were cultured onto plasma-treated PMMA with different membrane coating to model the extracellular microenvironment in vitro (Fig. 3B).

**Figure 3.**
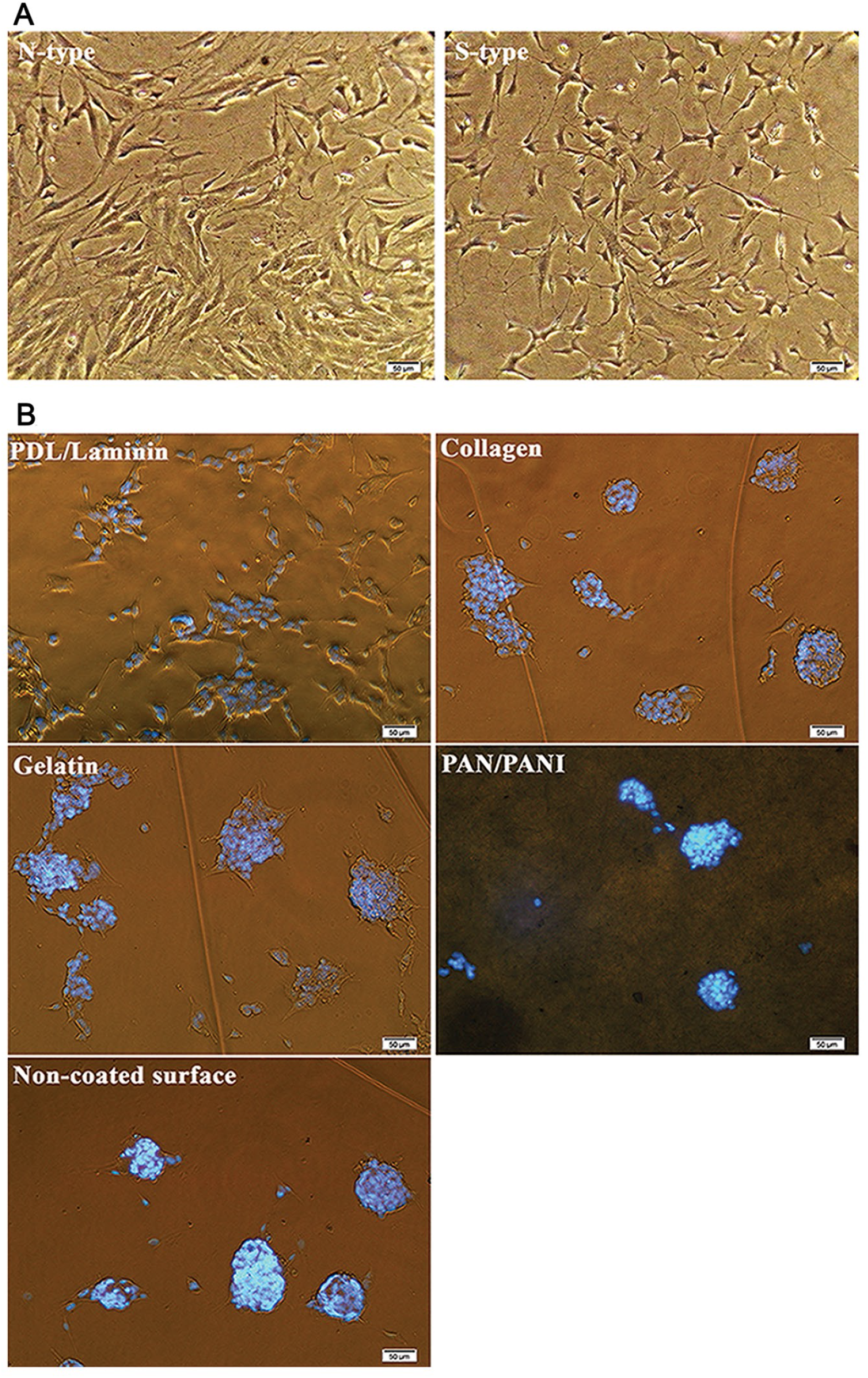
Phase contrast images of SH-SY5Y cells. Scale bars 50 μm. (A) Cells phenotypically changed from N-type to S-type by ATRA treatment. (B) Relatively greater number of differentiated cells oriented and aligned on different membranes compared to the naked surface as a negative control. PDL/laminin provided a physiologically optimal environment for neuronal adhesion and expansion.

The cellular density was significantly more ordered with PDL/laminin coating as compared to gelatin and collagen membranes, PAN/PANI nanocomposite or non-modified surface (negative control). The SH-SY5Y exhibited better differentiated morphology and more regular pattern of growth on the PDL/laminin membrane. Furthermore, the PDL/laminin membrane induced neurite elongation and branching, while other membranes increased cell aggregation, reminiscent of the aggregation commonly observed in undifferentiated cultures. Due to these reasons, we selected PDL/laminin surface coating for cell culture purposes in the following experiments.

### 3.3. Characterization of RGCs

The steps of RGC isolation by MACS technique are summarized in Fig. 4A. After isolation, flow cytometry showed high purity of RGCs, in which the percentage of RGCs was calculated by determining the ratio of CD90.1-positive cells (Fig. 4B). Isolation of RGCs was initially performed using a mesh filter according to RGC cell size (>40μm) to increase their population from a baseline of 19.8% in the single-cell retinal suspension to 37.4%. In the later stage MACS isolated RGCs according to CD90.1 surface marker in a ‘yes or no’ decision manner up to 70.4% purification. The RGCs exhibiting CD90.1 were also characterized using immunocytochemistry assay by above-mentioned antibodies and counterstaining with DAPI (Fig. 5).

**Figure 4.**
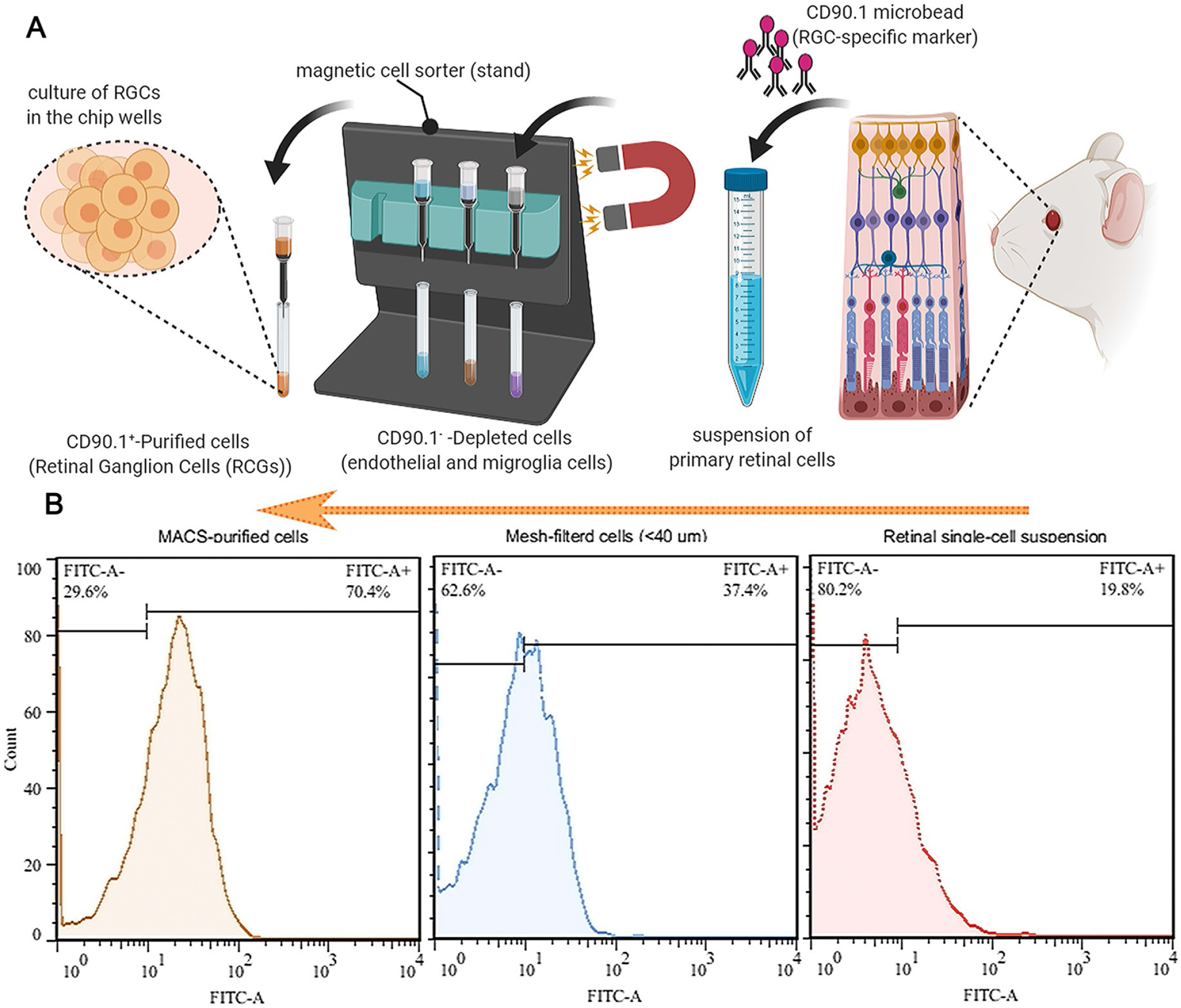
Purification of primary rat RGCs. (A) Retinal cells were isolated and converted into a cell suspension by triturating. The cell suspension was filtered through a mesh filter (>40 μm) to yield a single cell suspension. For magnetic assisted cell sorting (MACS), the single cell suspension was incubated with the RGC-specific marker CD90.1 microbeads and passed through an LS column to hold the micro bead-labelled cells (RGCs) in the magnetic field. The adherent RGCs were rinsed to deplete endothelial cells and microglia. The column was removed from the separator and the adhered cells were flushed out by firmly pushing the plunger into the column and counted. (B) Flow cytometry results showed that CD90.1+ cells were more purified after positive selection using MACS (orang, 70.4% purity) rather than mesh filtration (blue, 37.4%) and initial retinal single-cell suspension (red, 19.8%).

**Figure 5.**
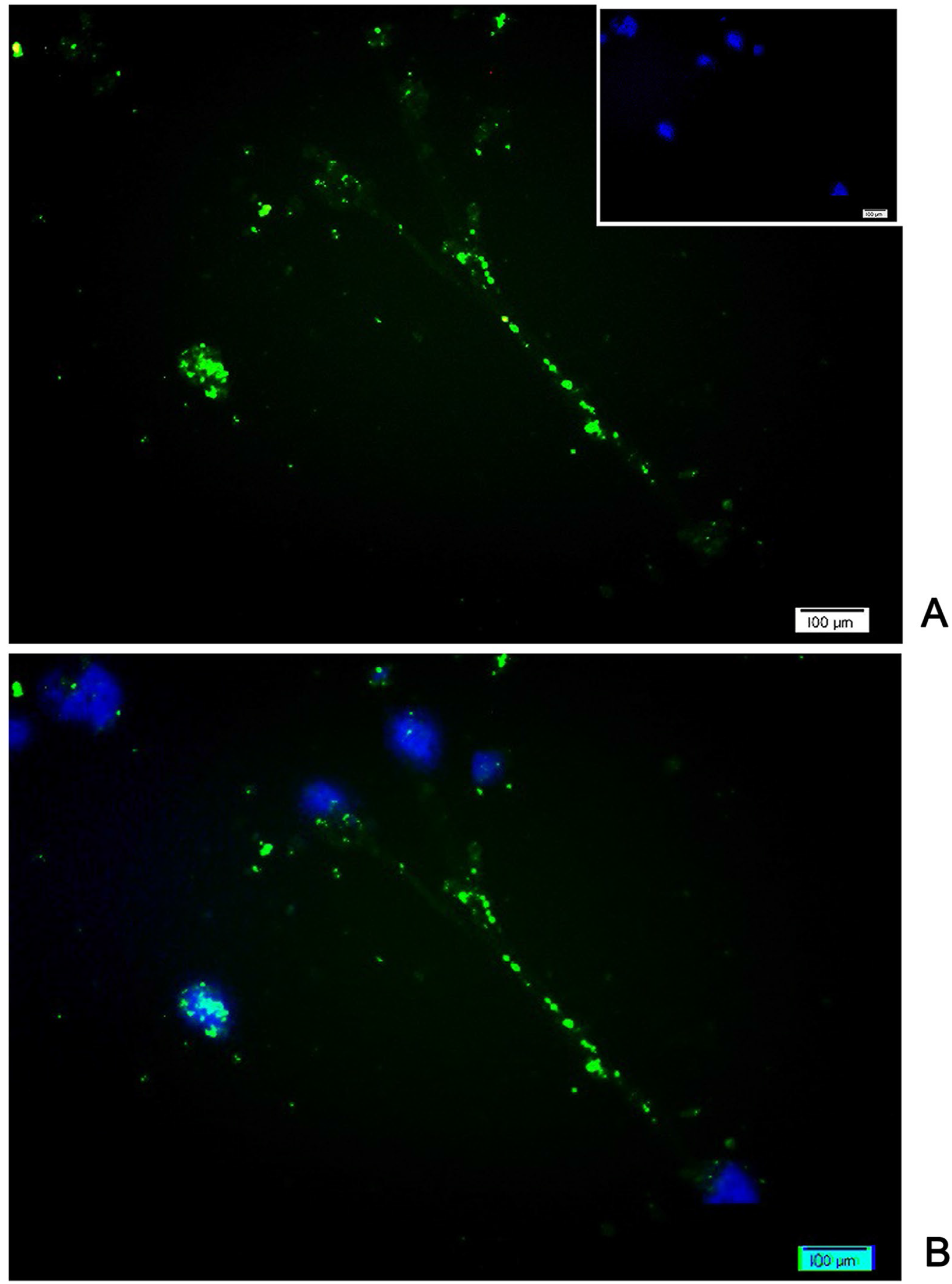
CD90.1 immunostaining of isolated rat RGCs by MACS. (A) After 3-day normal culture, RGCs continued to express the specific marker, CD90.1. Nuclear staining was performed using DAPI. (B) The axonal connections were observed between multiple RGCs in the merged image.

The RGCs were identified by their morphology and immunofluorescence staining characteristics. Dendrites and axons typically appear at opposing poles of the neuronal cell. On non-modified PMMA surface, RGCs tend to form cell clusters with small to media round cell bodies with few extended fine neurites after a 10-day culture. The majority of RGCs on the modified PMMA surface exhibited uniform and round cell bodies, with neurites extending to connect to one other. At day 10, the cells developed a complex dendritic network with numerous neurites and branches (Table 1).

**Table 1.**
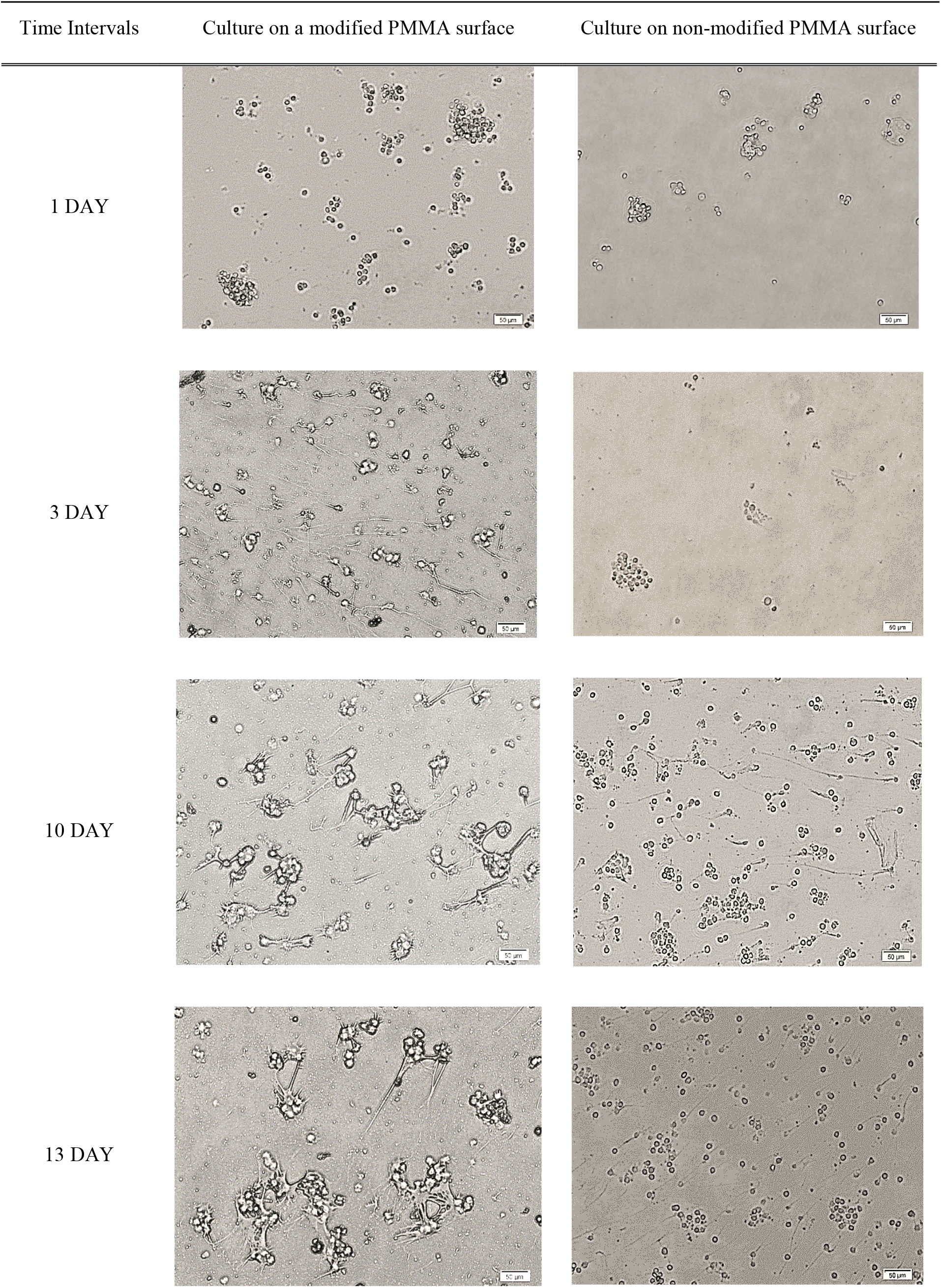
Morphological changes of primary RGCs at different time intervals on modified and non-modified PMMA surfaces before the pressure experiment. Scale bar= 50 μm.

### 3.4. Measurement of Cell Viability

One set of the chips was connected to a gas tank to apply elevated pressure at 33 mmHg, while another chip were exposed to normal pressure at 15 mmHg. As summarized in Figure 6, the EHP system consisted of a compressed 8% CO2/92% air gas tank, which was placed just outside the incubator and adjusted with both regulators for normal and ultra-low delivery pressure. Polyurethane hoses (4/8” ID) were used to feed the gas mixture into the pressure chip through a port at the back of the incubator. A digital pressure gauge (Ashcroft, DG25; 0.5% of 0-15psi span accuracy, ±0.1 mmHg sensitivity; relative to atmospheric pressure) was used to monitor the input pressure value. Pneumatic fittings were placed into the inlet/outlet circuits to provide tight control over gas flow. During loading, the pressure was also monitored by a sensor in the chip and kept at the 33 mm Hg with the accuracy of ±0.1 mmHg (data not shown).

**Figure 6.**
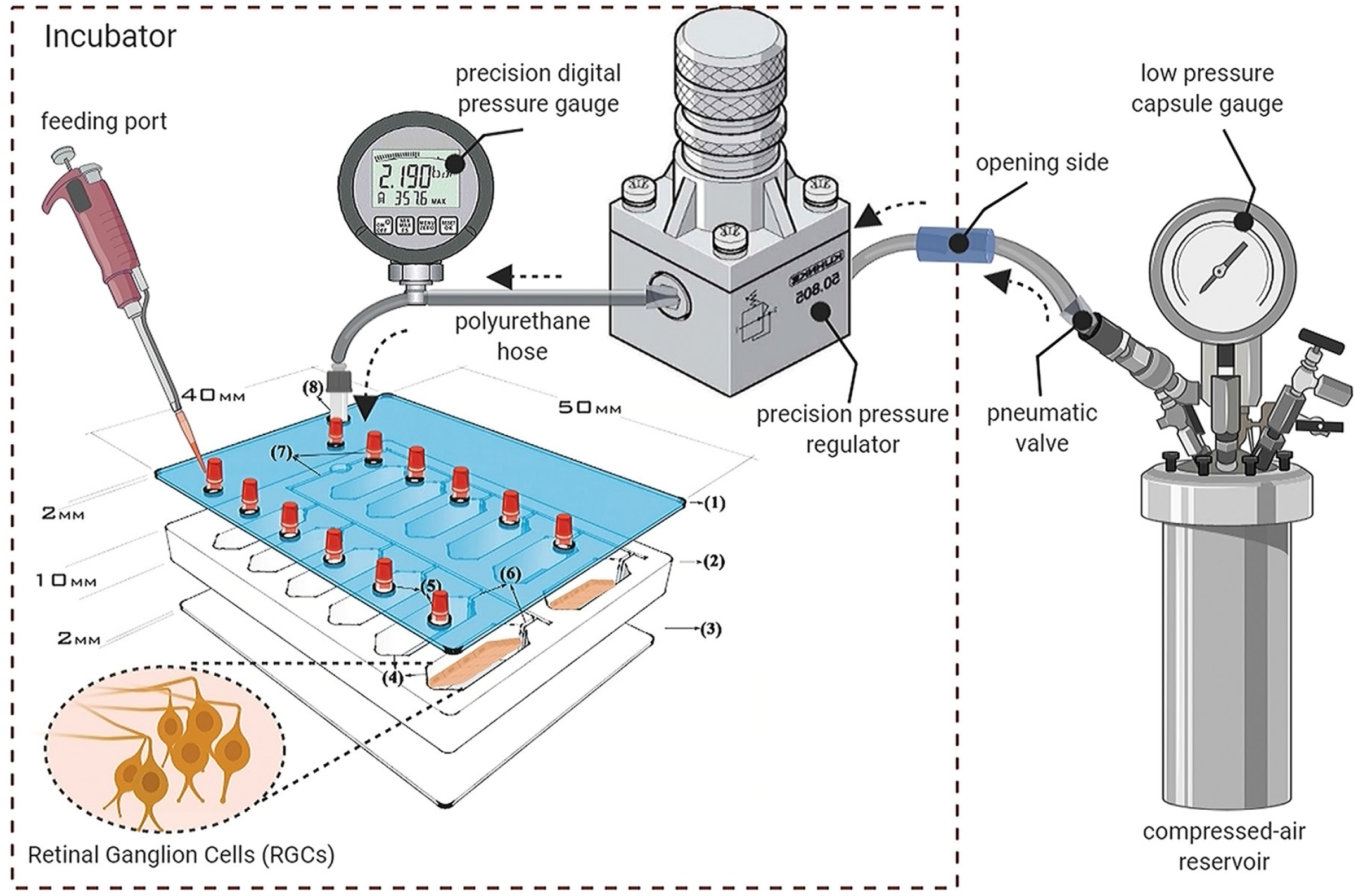
Schematic of the pressure set-up to simulate glaucomatous condition. A sealed pressure chip inside the incubator was connected to a gauge and regulators placed outside by the polyurethane hoses and pneumatic valves via a side opening in the back of incubator.

Untreated RGCs decreased in a time-dependent manner when elevated hydrostatic pressure was applied in the chip (Fig. 7A). From 6h of EHP onwards, a significant difference was observed in cell survival as compared to control cells under normal hydrostatic pressure. Cell viability results showed 85, 78, 70, 67 and 61 percent cell survival under normal pressure versus 40, 22, 18, 12 and 10 percent cell survival under high pressure at 6, 12, 24, 36 and 48 hours (p <0.0001), respectively. BDNF-treated RGCs demonstrated better cell survival with or without elevated pressure as compared to untreated cells. BDNF-treated RGCs showed 92, 89, 88, 83 and 80 percent viability under normal pressure compared to 72, 67, 64, 60 and 53 percent under high pressure at 6 (p =0.0198), 12 (p =0.0022), 24 (p =0.0004), 36 (p <0.0001) and 48 (p <0.0001) hours, respectively (Fig. 7B). RNYK provided slightly more stability than BDNF over time. RNYK-treated RGCs showed 94, 93, 89, 87 and 86 percent viability under normal pressure compared to 67, 66, 63, 60 and 58 percent under high pressure at 6, 12, 24, 36 and 48 hours (p <0.0001), respectively (Fig. 7C). BDNF and RNYK treatments induced separately an approximate two-fold decrease in RGC death rates as compared to untreated cells under both normal and elevated pressure condition.

**Figure 7.**
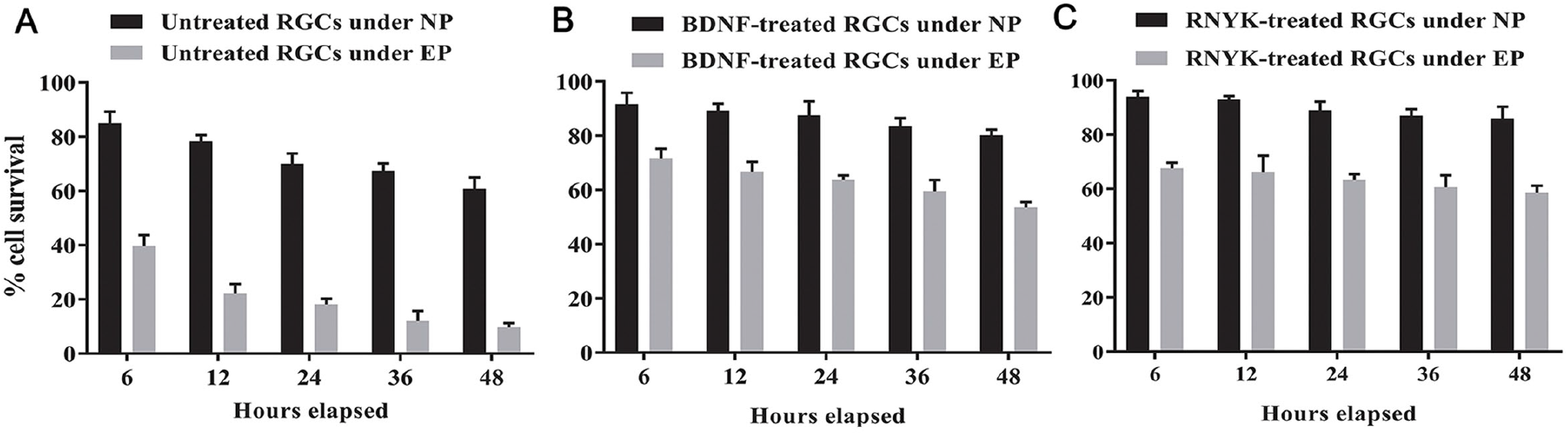
Cell survival assay under NP* and EP** conditions (Two-way ANOVA, N=4, graphs display mean ±SD). (A) There was a significant decrease over time for untreated cells under EP condition with mean difference (Mean Diff.) =45.35, 56, 51.83, 55.43, and 51.2 at 6, 12, 24, 36 and 48 hours, respectively (p<0.0001). (B) BDNF prevented degeneration of treated RGCs under EP condition with Mean Diff. = 20, 22.52, 23.83, 24, and 26.7 at 6, 12, 24, 36 and 48 hours, respectively (p<0.0001). (C) RNYK identically protected RGCs under EP condition with Mean Diff. = 26.33, 26.67, 25.67, 26.33, and 27.33 at 6, 12, 24, 36 and 48 hours, respectively (p<0.0001).

## 4. Discussion

Pressure chambers are important tools to study the influence of elevated pressure on individual cell types [38]. Prior to employing animal models, they allow the assessment of pre-defined experimental conditions and evaluation of specific variables contributing to cell survival. Over the past two decades, various studies have reported significant changes in the behaviour of different cells following the application of hydrostatic pressure [22, 26, 39–47]. These changes include increased apoptosis, changes in cell morphology and gene expression. Neuronal cells including PC12 [48, 49] and RGC-5 cell lines [23, 50–53], primary cultures of RGC [54, 55], optic nerve head astrocytes [24, 27, 28, 56–61], Müller cells [62–64], or microglial cells [65–67], and more complex preparations such as organotypic co-cultures of the retinal cells [25, 68, 69] or eye cup preparations [15, 17, 70, 71] have been exposed to elevated pressure.

In the vast majority of glaucoma patients, IOP levels may vary from 20 to 35 mmHg. Several in vitro glaucoma models have been developed to improve the accuracy and repeatability of experimental conditions in this pressure range. Pressurized chamber is the most commonly used model established by Kosnosky et al. (1995) and modified by Agar at al. (2000) to investigate cellular mechanisms [49, 72]. In this system, a mixture of humidified gas is injected through a pressure regulator into a glass macro-chamber with ±1.5 mmHg variation. Studies using this chamber have indicated that exposure of RGC-5 cells to 30, 50, or 100 mmHg for 72, 6, or 2 h, respectively, triggers apoptosis via oxidative stress and mitochondrial dysfunction, similar to the human condition when comparing acute glaucoma (100 mmHg) with chronic glaucoma (30 mmHg) [23, 50–52, 73–75]. The second important model is a hydraulic pressurizing chamber introduced by Parkkinen et al. (1993), which compressed the internal gas phase using a hydraulic system [76]. Tezel and wax (2000) used this device to expose RGCs to 50 mmHg for 72 h and promote neurodegeneration [22, 25].

In the present study, we designed a glaucoma-on-a-chip system that provides continuous hydrostatic pressure with ±0.1 mmHg variation under a temperature-controlled environment. Our system was custom-made in agree with the pressurized chamber approach, thus, several conditions and confounding factors were carefully simulated, validated or controlled beforehand. They consisted of fluid velocity fields, alterations in gas composition, temperature and osmolarity to design a more stable and adjustable experiment compared to prior in vitro studies. Our results indicated that glaucoma-on-a-chip can simulate glaucomatous conditions only when provided with a suitable microenvironment for cell-to-cell communication of RGCs. Accordingly, the PMMA surface was modified by air plasma and a thin layer of PDL/laminin to form a continuous surface of mechanical properties and uniform chemistry leading to a homogenous cell distribution. Furthermore, the evaluation of cellular mechanisms requires isolation and culture of primary RGCs. Early postnatal tissues were used to optimize RGC viability in primary cell cultures due to the limited yield and the typically post-mitotic feature of these neurons in the next state. The use of RGCs in cultures on chips allowed examining a direct in vitro situation in which retinal cells maintain their microarchitecture.

Most reports investigating the degenerative effect of chronic IOP elevation have used pressures ranging from 30 mmHg (for 72 h) to 100 mmHg (for 2 h), as already described. In the present study, we exposed RGCs to 33 mmHg (elevated hydrostatic pressure, EHP) or 15 mmHg (normal pressure) for 6, 12, 24, 36 and 48 hours, with and without adding neuroprotective agents. Accordingly, cell survival of primary rat RGCs was evaluated under four different experimental conditions simulating the high pressure glaucoma (untreated RGCs under EHP), normal-tension glaucoma (BDNF deprived RGCs under normal pressure), neuroprotective-treated glaucoma (BDNF/RNYK treated RGCs under EHP), and normal conditions (BDNF-treated RGCs under normal pressure). Our results demonstrated that BDNF can maintain neuronal viability despite high pressure. According to cell survival patterns and rates, the neuroprotective influence of BDNF seems to cancel the detrimental effect of elevated pressure. This suggests that the negative impact of neurotrophin deprivation can be compared to the insult of high pressure. This may be equivalent to the observation that failure of axonal transport of neurotrophins in glaucoma triggers RGC degeneration and that retinal cells essentially require neurotrophic input from target neurons in the brain.[77, 78]

This previously confirmed that RNYK can mimic BDNF activities in vitro at an optimum concentration of 5 ng ml^−1^. RNYK prevented neuronal degeneration of ATRA-treated SH-SY5Y with equal efficacy to or even better than BDNF at 50 ng ml^−1^ [33]. In the current study, RNYK indicated a neuroprotective effect comparable to BDNF on primary RGCs under both normal and elevated pressures. In light of the findings obtained, the identified peptidomimetic compound is significant as a promising neuroprotective agent for glaucoma treatment. Also, small size of RNYK might be an advantage for drug design and synthesis in future. It is clear that in vitro models may never substitute animal studies, but they are important tools in preclinical studies. Glaucoma-on-a-chip offers the advantages of allowing controlled experimental conditions, preliminary targeting of a specific cell type or pathway involved in glaucomatous damage. More studies are needed to develop models to study RGC neurodegeneration and neuroprotection by putative agents such as neurotrophins, peptides, and other small molecules prior to assessment in animal models.

## Acknowledgements

We are thankful to Dr. S Gharavi (Alzahra University, Tehran, Iran) and Dr. N K Tafreshi (Moffitt Cancer Center and Research Institute, Florida, USA) for technical support.

